# Targeted Irradiation and STAT3 Inhibition Reprogram the AML Microenvironment and Extend Survival: Toward Translational Immunoradiotherapy

**DOI:** 10.1101/2025.05.14.653587

**Authors:** Srideshikan Sargur Madabushi, Jamison Brooks, Darren Zuro, Yu-Lin Su, Damian Kaniowski, Hemendra Ghimire, Ji Eun Lim, Amr Mohamed Hamed Abdelhamid, Ya-Huei Kuo, Guido Marcucci, Jeffrey Wong, Monzr Al Malki, Anthony Stein, Marcin Kortylewski, Susanta Hui

## Abstract

**Purpose:** Despite advances in acute myeloid leukemia (AML) therapy, relapses remain challenging. While AML is radiation-sensitive, total body irradiation (TBI) causes organ toxicities and activates tolerogenic/proangiogenic STAT3 signaling. CSI-2, a myeloid cell-targeted STAT3 inhibitor, promotes anti-leukemic immune responses but has limited efficacy against high disease burden. We investigated whether image-guided targeted marrow irradiation (TMI), which focuses radiation on leukemia sites while sparing critical organs, could synergize with CSI-2 to improve leukemia clearance and establish durable immunity.

**Methods:** Mice were intravenously engrafted with CMM-AML cells reaching 20–30% in bone marrow (BM) infiltration (moderate-to-high disease burden) before receiving IV-injections of CSI-2 (5mg/kg) with or without TMI. Fluorescently labeled CSI-2 biodistribution was assessed using flow cytometry and quantitative multiphoton microscopy. Survival was monitored for 3–4 months before evaluating BM composition using flow cytometry and immunohistochemistry.

**Results:** TMI significantly improved vascular permeability and scavenger receptor/TLR9-dependent uptake of CSI-2 by AML cells and leukemia-associated myeloid cells. Combined TMI/CSI-2 treatment more effectively reduced high leukemia burden than CSI-2 alone, achieving >80% survival at 120 days with increased CD8+ cytotoxic and CD4+ helper T cell infiltration. TMI/CSI-2-treated mice were protected from AML rechallenge suggesting that they developed protective immune memory. In an aggressive MLL-AF9 AML model, TMI/CSI-2 combination significantly extended survival compared to either monotherapy.

**Conclusion:** TMI/CSI-2 strategy represents a novel organ-sparing immunoradiotherapy that synergistically enhances leukemia clearance while promoting long-term protective immunity. These findings warrant further investigation of this strategy for high-burden or relapsed AML and provide the foundation for clinical translation.

## Introduction

AML is one of the most common types of leukemia, representing 38.4% (20,830) of all leukemia cases and 42.8% (10,460) of leukemia-related deaths.(1) Despite advances in chemotherapy treatment and allogeneic hematopoietic stem cell transplantation (allo-HCT), only 40% of younger (<60 years) patients and 10% of older patients (>60 years) survive. Relapsed/recurrent AML carries a dismal 5-year overall survival of 7-20%; thus, there is an urgent and unmet need for more effective therapies.(2-6)

High-intensity ionizing radiation has the potential to reduce leukemia burden.(7) However, toxicities related to high-dose total body irradiation (TBI) offset any gains in overall survival.(8) To address this, we developed a computed tomography-based, image-guided intensity modulated total marrow irradiation (TMI) platform. TMI selectively target radiation to the disease-rich sites such as the bone marrow and lymphoid tissues, while sparing vital organs (e.g., lungs, heart, liver). Using this platform, we escalated bone marrow (BM) doses to 2000 cGy (2 Gy × 10 fractions) in combination with cyclophosphamide and etoposide as conditioning for allo-HCT in patients with advanced, relapsed, or refractory AML.(9) In a Phase II trial (NCT#02094794) involving 74 patients, TMI-based conditioning achieved a 2-year progression-free survival (PFS) 33%, overall survival (OS) 46%, and cumulative incidence of non-relapse mortality (NRM) of just 12% (10) representing a substantial improvement over historical 2-year OS rates (< 10%) with conventional TBI regimens(11). Nevertheless, more than half of the patients experienced relapse within one year, underscoring the urgent need for enhanced, combinatorial therapeutic strategies.

To enable mechanistic and therapeutic discovery in a preclinical setting, we developed a clinically analogous, image-guided TMI platform for use in murine models of AML.(12) This platform faithfully mimics human TMI, enabling organ-sparing radiation delivery with high spatial precision, and has been shown to minimize off-target tissue injury in preclinical models.(13)The ability to recapitulate clinical TMI in mice allowed us to reverse translate key clinical questions into mechanistic studies and therapeutic testing. In reverse translation to further explore new treatment development, we developed clinically equivalent image guided targeted TMI in murine model [Zuro et al] and show sparing critical organs, prevent tissue damage.(13) These developments further allowed us to implement TMI to investigate mice with AML model.

The principles of cancer radiotherapy are based on the generation of direct cytotoxic effects to tumor cells with a potential to trigger systemic antitumor immune responses. (14) However, such beneficial, abscopal immune effects are clinically rare. We previously demonstrated that cancer irradiation is more likely to stimulate inflammatory signaling promoting tumor vascularization and regrowth. (15) DNA released by dying tumor cells induces Toll-like receptor-9 (TLR9) activation in myeloid cells, such as macrophages, leading to interleukin-6 (IL-6) production and subsequent activation of STAT3, an oncogenic transcription factor and central immune checkpoint regulator.(15, 16) STAT3 promotes immune evasion by upregulating immunosuppressive molecules such as PD-L1 and arginase-1 (17, 18) (19), and by reducing MHC class II expression on antigen-presenting cells.(20, 21) Constitutive STAT3 activation is also commonly observed in AML blasts and correlates with poor prognosis.(22, 23) Dual role of STAT3, in promoting leukemic cell survival and immune evasion, make it an attractive target for AML therapy.(23-26) Notably, STAT3 inhibition does not affect the viability of normal immune cells,(15, 27) and dominant-negative mutations in STAT3 are not lethal in humans.(28, 29) suggesting a favorable therapeutic index. However, because STAT3 lacks enzymatic activity, it has proven difficult to inhibit with small molecules (30, 31). The initial oligonucleotide-based strategies to STAT3 inhibition, such as antisense oligonucleotides[28] or decoy oligodeoxynucleotides (dODNs) (30, 32) were hindered by inefficient and non-selective delivery.(33, 34)

To overcome delivery challenges, we developed a CpG-based delivery platform that leverages synthetic TLR9 agonists for myeloid cell-selective delivery of oligonucleotide therapeutics, such as siRNAs (35, 36), miRNA (37) or dODNs (38). CpG-conjugates target TLR9-expressing immune cells, including leukemia stem and progenitor populations, in vitro and in vivo. (36, 39, 40) The second generation CpG-STAT3 inhibitor (CSI-2) based on nuclease-resistant decoy DNA allows for rapid inhibition of STAT3 in human and mouse TLR9^+^ AML blasts and normal myeloid cells [PMID: 26796361]. CSI-2 demonstrated long-term antitumor efficacy against both human and mouse AML in vivo, by enhancing the immunogenicity of leukemic cells and promoting long-term antitumor T-cell immune responses.(41) While CSI-2 proved effective against lower-to-moderate AML burden, the high disease burden and rapid progression limits time required for T cell– mediated immunity to develop. Thus, we hypothesized that combining CSI-2 with organ-sparing TMI could rapidly reduce disease burden while priming the immune system, overcoming TLR9– STAT3–mediated tolerance and enabling long-lasting antitumor immunity. In this study, we leveraged this novel preclinical platform to investigate the therapeutic and immunologic potential of combining targeted low-dose TMI with CSI-2, a CpG-based STAT3 inhibitor, as a treatment for high-burden AML models in mice.

## Materials and Methods

### Human BM samples

This phase 2 clinical trial was registered with clinicaltrials.gov (NCT02094794) and approved by the City of Hope Institutional Review Board (IRB #14012). Informed consent was obtained for all study participants in compliance with the Declaration of Helsinki.

### Animal studies

All animal experiments were conducted in accordance with established guidelines from the Institutional Animal Care and Use Committee (IACUC) at City of Hope (COH). C57BL/6 mice, both male and female, aged 8-10 weeks were obtained from the National Cancer Institute (NCI). TLR9-KO/C57BL/6 mice were procured from Jackson laboratory (#034449) and maintained by animal breeding core at City of Hope. The CMM cells (Cbfb-MYH11/Mpl+) were established as described in prior studies. (38, 42) The CMM cells were expanded in mouse and spleen cells were harvested after 2-3 weeks from primary mouse and used for studies. MLL-AF9 AML cells were generated as previously described. (43) MLL-AF9 cells are established in vitro cell line and expanded in regular RPMI 1640 supplemented with 10% FBS 1% Glutamax, 1% Penicillin/streptomycin media and 1X β-mercaptoethanol (55μM). Cells were maintained in culture for 2 weeks before use and used within 4 weeks in culture. Both CMM and MLL-AF9 cells carry GFP reporter gene and is used to determine disease burden by flow cytometry.

### Mice Model

C57BL/6 or TLR9KO/C57BL/6 mice were injected via the lateral tail vein with 1 × 106 CMM-GFP AML cells suspended in phosphate-buffered saline (PBS), respectively. The percentage of circulating green fluorescent protein positive (GFP^+^) CMM cells was monitored by flow cytometry, and once it was between 5% to 10%, the mice were treated six times daily with RO containing CSI2 (5 mg/kg) or PBS. The first dose of treatment, with or without RO, was combined with TMI at a dose of 4 Gy using the Precision X-RAD SMART Plus/225cx irradiator (Precision X-Ray, CT, USA).

### AML disease kinetic study

Mice were injected with CMM (1 million GFP+ cells) cells via tail vein/ retro orbital. GFP+ cells in blood and bone marrow were assessed from Day 4-, 6-, 8- and 12-days post injection, n=3/timepoint. Blood to BM correlation was plotted and 5-10% blood corresponds to 20-30% in Bone marrow and was used as intervention disease burden in this study. When the disease burden was 5-10%, CMM bearing mice were intervened ± TMI/ CSI-2. CSI-2 (5mg/Kg) was injected every other day (via RO), starting Day 0, 30 min-1h prior to TMI treatment for 6 doses. TMI treated was designed and delivered as previously described. PBS treated mice was used as control. Same QODX6 dosing was carried out for MLL-AF9 intervention post TMI.

### Oligonucleotide design and synthesis

CpG oligonucleotides and their conjugates were synthesized at the DNA/RNA Synthesis Core (COH) by linking CpG-1668 ODNs to STAT3 decoys, following previously described methods. (44)

### Flow cytometry

Single-cell suspensions from mouse spleen and bone marrow were prepared by enzymatic and mechanical dissociation, while leukocytes were isolated from peripheral blood as previously described. (45) Surface staining was performed using fluorochrome-conjugated antibodies against CD11b, CD8, CD4, CD44, CD62L, CD19, Gr-1, F4/80, MHC-II and CD86, following Aqua LIVE/DEAD staining and FcγIII/IIR blockade. Intracellular staining was carried out using the Fix/Perm kit with antibodies targeting, IFNg, FoxP3 and pSTAT3. Fluorescence data were acquired using BD LSRFortessa, NovoCyte Quanteon (Agilent), flow cytometers and analyzed with FlowJo v10 software (TreeStar, Ashland, OR).

### Transcriptomic profiling of patient BM samples using Nanostring

Differential gene expression in Human patient samples, relapse (n=7) and remission (n=5) were carried out using he Nanostring Human PanCancer Immune Panel. RNA was extracted using miRNeasy FFPE kit (Qiagen cat# 217504), RNA concentration was assessed with the Nanodrop spectrophotometer ND-1000 and Qubit 3.0 Fluorometer (Thermo Scientific, CA). RNA fragmentation and quality control were determined by 2100 Bioanalyzer (Agilent, CA). RNA expression was analyzed by NanoString nCounter platform (NanoString Technologies, WA) using Human PanCancer Immune Profiling panel consisting of 770 genes. RNA was first hybridized with Codeset from gene panel at 65°C for 16 hours. Post-hybridization probe-target mixture was the purified and quantified with nCounter Digital Analyzer.

All raw data from expression analysis were first aligned with internal positive and negative controls, then normalized to the selected housekeeping genes included in the assay. Differential gene expression patterns as well as pathway scores with statistical analyses were performed with nSolver software (NanoString Technologies, WA). Pathway Scores were calculated by nSolver Advance Analysis module to summarize the data from a pathway’s genes into a single score, using the first principal component (PC) of the expression data (46). In brief, PC analysis scores each sample using a linear combination of its gene expression values, weighting specific genes to capture the greatest possible variability in the data. The data are presented in a log2 scale, in general, increased score indicates increased overall expression.

### Immunohistochemistry

Histology (H&E) and Immunohistochemistry for CMM-GFP, CD4 and CD8 was carried out according to SOP at City of Hope pathology core. Details in supplementary methods section.

### Statistics

Statistical significance between two experimental groups was determined using an unpaired two-tailed t-test. For comparisons among multiple treatment groups, one-way ANOVA followed by Bonferroni post hoc testing was performed. P values, with significance denoted as follows: ***P < .001; *P < .01; P < .05. All statistical analyses were conducted using Prism software (version 9.5; GraphPad).

All other materials and methods are provided in supplementary sections.

## Results

### STAT3 activity in AML-associated macrophages is reduced in patients sensitive to radiotherapy but not in patients with disease relapse

The ongoing phase II trial (NCT#02094794) to assess the safety and efficacy of total marrow and lymphoid irradiation (TMLI) in combination with chemotherapy drugs before donor stem cell transplant indicated potential benefits to patients with AML.(9) However, despite improvements in the overall and progression-free survival more than half of the patients experienced relapse within one year, underscoring the still unmet need for further optimization of such combinatorial radiotherapeutic strategy. STAT3 activation is known to underscore AML survival and therapeutic resistance, thus we assessed whether it plays a role in the AML patients’ sensitivity to TMLI treatment. We analyzed STAT3 activation (tyrosine 705 phosphorylated pSTAT3) in bone marrow (BM) biopsies derived from AML patients responding to TMLI therapy compared to patients with confirmed AML relapse. The multiplex IHC analysis indicated that in treatment-sensitive patients pSTAT3 levels decreased in both leukemia-associated macrophages (CD163^+^) and in CD34^+^ stem cells, potentially leukemia stem cells (LSC) (**Fig 1A**), although the reduction was statistically significant only in macrophages (**Fig 1B-C**). In contrast, the patients with AML relapse retained high STAT3 activation in both cell populations. As shown before, (PMID: 26796361) TLR9 was consistently and commonly expressed in most CD34^+^ AML cells and in leukemia-associated macrophages both pre- and post-TMI-based transplant conditioning regimen for allogeneic HCT (**Fig 1A**). Importantly, STAT3 activity remained reduced in treatment-sensitive patients even after 100 days from the treatment (**Fig 1D**). These observations suggested that targeting STAT3 signaling in TLR9^+^ leukemia-associated macrophages and in AML cells could interfere with the potential mechanism of leukemia relapse after TMLI.

**Figure 1.**
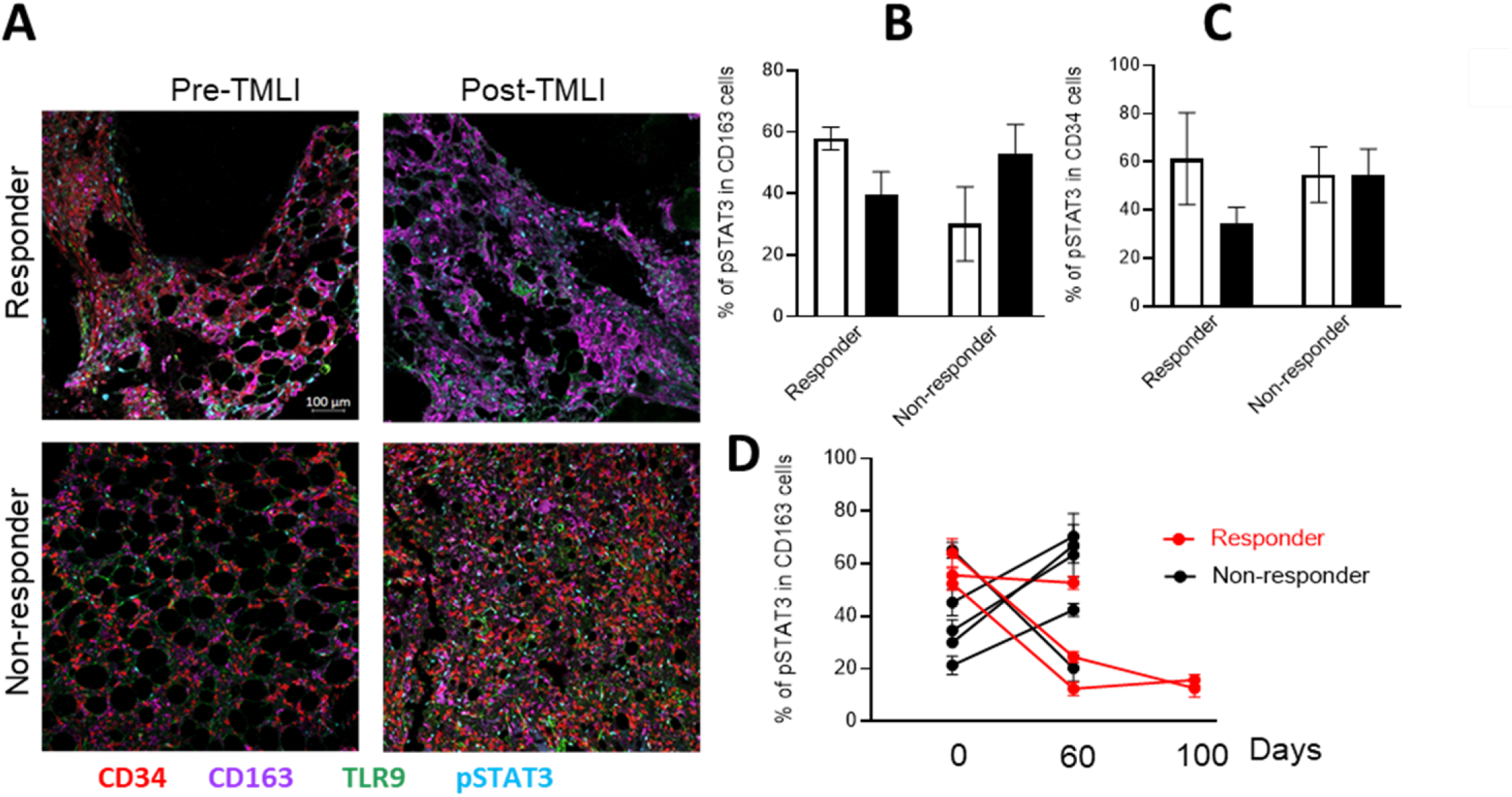
STAT3 activity in leukemia-associated macrophages is reduced in AML patients sensitive to radiotherapy but not in patients with disease relapse. **(A-C)** STAT3 activation (blue) and TLR9 expression (green) in leukemia-associated CD163+ macrophages (magenta) or in CD34+ cells representing primarily AML cells (red) in bone marrow biopsies from AML patients responding and non-responding to TMLI before and after 60 days from the treatment. Representative images of the multiplex IHC staining (Perkin-Elmer) **(A)** and quantification (QuPath sofware) of the percentage of pSTAT3+ cells **(B-C). (D)** The reduced STAT3 activity in leukemia-associated macrophages is long-lasting and detectable after 100 days from treatment. **(D)** after Shown are means±SD (non-responding AML patients, *n*=5; treatment-sensitive AML patients, *n*=3).

### Targeted marrow irradiation enhances blood flow and CSI2 oligonucleotide uptake by target AML cells in vivo

Our initial studies determined the *Cbfb/MYH11/Mpl* (CMM) leukemia progression kinetics in C57BL/6 mice. As shown in Figure 2A, there was a correlation between levels of CMM cells in peripheral blood and in bone marrow with ∼5-10% of AML cells in blood corresponding to ∼25-30% of BM-localized AML cells (**Fig. 2A**). CMM leukemia model was previously described to have high levels of STAT3 activity in leukemic cells and in the AML microenvironment (38, 41) similarly to patients with recurrent AML (**Fig. 1**). To disrupt tolerogenic STAT3 signaling in CMM microenvironment and enhance sensitivity to radiation therapy, we combined TMI with an oligonucleotide strategy inhibiting STAT3 specifically in TLR9-positive myeloid cells, CpG-STAT3decoy or CSI2. (41) CMM-bearing mice (5-10% of AML cells detectable in peripheral blood) were treated using intravenous injection of 5 mg/kg CSI2 followed by a single dose of TMI. To reduce the uncertainty about the temporal dynamics of radiation effects,(47) we employed quantitative multiphoton microscopy (QMPM) for longitudinal imaging of AML progression and vascular function in mice with high leukemia burden (≥30% in BM) through a cranial window 2-5 days after TMI (**Fig. 2B**). Initial studies in CMM-AML-bearing mice treated using a single 2 Gy dose of TMI showed improved BM vascular parameters such as blood flow and vessel diameter (**Fig.2C**) and vascular permeability as measured by K_trans_ (**Fig. 2D**). Notably, low dose 2 Gy TMI restored vascular permeability to baseline levels more rapidly than 10 Gy, suggesting a dose dependent relationship between TMI and vascular recovery. Next, we compared the effect of two doses of TMI, 2 and 4 Gy, on leukemia clearance. As expected, 4 Gy TMI dose more significantly reduced leukemia burden as indicated by fewer percentage of GFP^+^ CMM cells in the BM and decreased splenomegaly compared to the 2 Gy group (**Supplemental Fig. S1**).

**Figure 2.**
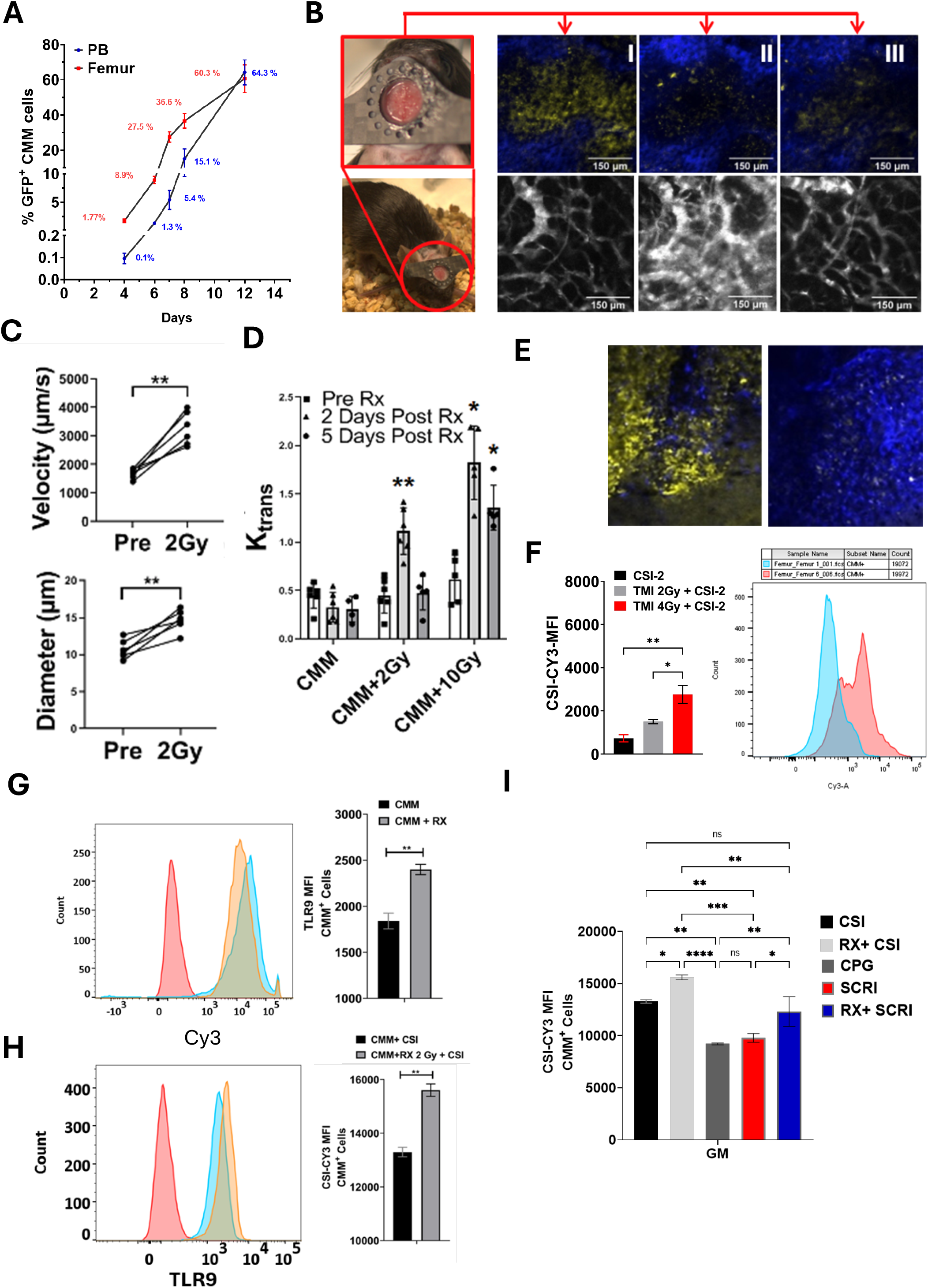
Targeted marrow irradiation enhances blood flow and CSI2 oligonucleotide uptake by target AML cells in vivo. C57BL/6 mice was engrafted with GFP-expressing CMM leukemia cells. **(A)** The leukemia progression was assessed in PB and bone marrow (femurs) using flow cytometry at different time points; means±SEM (n=3). **(B-D)** Radiation augments CSI-2 delivery and increases uptake by CMM cells. (**B)** QMPM imaging of mouse calvarium using cranial window. **(C-D)** TMI (2 Gy) increased rate of blood flow and vessel diameter **(C). (D)** The vessel leakiness is TMI dose-dependent and reversible. **(E, F)** TMI enhances CSI2 uptake by leukemic cells in vivo. QMPM imaging **(E)** and flow cytometric analysis **(F). (G, H)** Irradiation (2 Gy) upregulates CSI2 uptake **(G)** and TLR9 levels **(H)** in cultured CMM cells; means±SEM (n=3). **(I)** CSI2 uptake is scavenger receptor-dependnent; SCRI: scavenger receptor inhibitor=dextran sulfate; CpG: unlabeled CpG oligonucleotide competitor.

To assess whether these vascular changes translated to improved oligonucleotide delivery, we evaluated CSI2 uptake in CMM-bearing mice treated with or without TMI (2 or 4 Gy) (**Fig. 2E-H**). Mice received fluorescently labeled CSI2^Cy3^ 48 h after TMI, and the oligonucleotide uptake by CMM cells was analyzed using in vivo QMPM imaging (**Fig. 2E**) and flow cytometry (**Fig. 2F**). The imaging through a cranial window confirmed increased CSI-2 accumulation in CMM cells after TMI (**Fig. 2E**) and correlated with the significantly enhanced CSI2 uptake by leukemic cells compared to CSI2 alone, especially after 4 Gy irradiation (**Fig. 2F**). Together, these data suggested that TMI enhances both CSI2 delivery and uptake, likely through improved vascular perfusion and permeability. Based on these findings, we selected the dose of 4 Gy TMI, that showed improved leukemia cell killing and improved CSI2 delivery, for use in all subsequent TMI-CSI2 combination experiments.

To further characterize how radiation affects CSI-2 uptake in leukemic cells, we conducted in vitro assays using CMM cells treated with 2 Gy radiation. One day after irradiation, cells showed increased uptake of CSI2^Cy3^ as assessed by flow cytometry (**Fig. 2G**). Radiation also upregulated TLR9 expression in CMM cells that could result in better endosomal escape of the oligonucleotide and improved immunostimulatory effects (**Fig 2H**).(PMID: 23777886) Importantly, the augmented oligonucleotide uptake by CMM cells after irradiation was abrogated by an unlabeled CpG oligonucleotide (TLR9 agonist) as well as by dextran sulfate used as a scavenger receptor Inhibitor (**Fig. 2I**), suggesting the role of scavenger receptors in the internalization of CSI2 similarly to other CpG-containing oligonucleotides.

### TMI/CSI2 combination treatment promotes T-cell recruitment and leukemia regression

We have compared the efficacy of TMI and CSI2 as single agents and in combination using syngeneic CMM and MLL-AF9 leukemia models representing two different genetic types of AML, inv(16) and t(9;11)(p22;q23) translocation related leukemias. As shown in **Fig. 3A**, the combination TMI/CSI2 treatment improved mice survival in both tested AML types compared to limited efficacy of either treatment alone. In mice with moderate-to-high CMM, the TMI-CSI-2 combination resulted in long-term disease control, with >80% of treated mice surviving beyond 120 post-intervention. In contrast, CSI2 monotherapy modestly extended median survival to 48 days, while TMI alone produced only a slight improvement (∼10.5 days) compared to untreated controls. Similar effect was observed in MLL-AF9 model (**Fig. 3A**, right panel). TMI/CSI2 co-treatment led to a median survival of ∼82 days, with ∼50% of mice achieving disease-free survival. In contrast, TMI alone (median survival ∼31 days) and CSI-2 alone (median ∼10 days) had limited therapeutic benefit compared to untreated controls (median ∼8 days). These results indicate that TMI/CSI2 combination can be an effective immunotherapeutic strategy not limited to a single subtype of AML.

**Figure 3.**
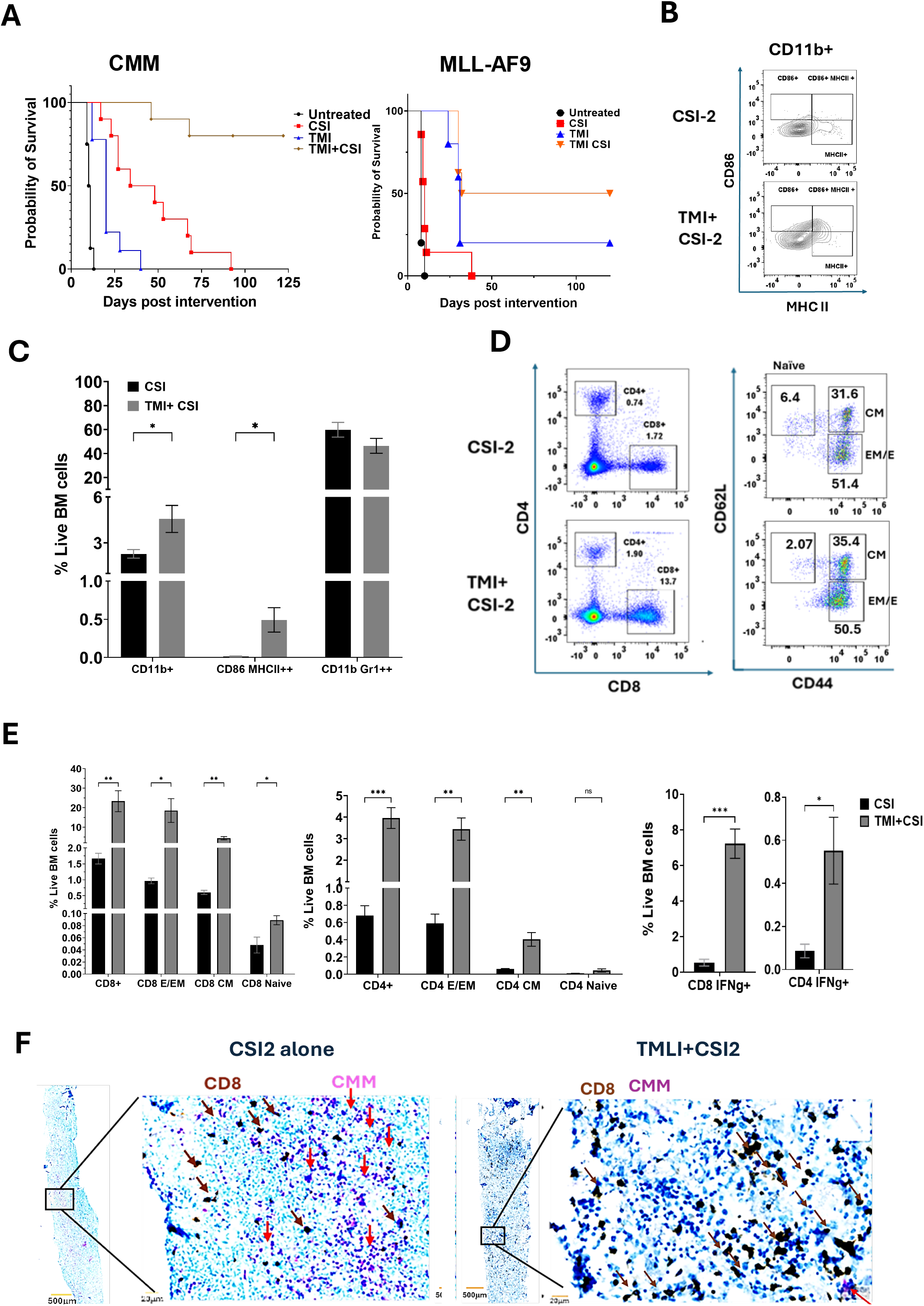
TMI/CSI2 combination treatment potently activates leukemia-associated myeloid cells, resulting in T-cell recruitment and leukemia regression in mice. **(A)** The combined TMI/CSI2 treatment resulted in long-term animal survival of mice engrafted with two different types of AML, CMM (keft panel) or MLL_AF9 (right panel). Combined results derived from two independent studies; means ± SEM (n=8). **(B-C)** TMI/CSI2 treatment activates antigen-presentation in bone marrow myeloid cells while reducing the percentage of tolerogenic MDSCs (CD11b+/Gr1+). Representative flow cytometric analysis **(B)** and the quantification of results **(C)**; means±SEM (n=3-6). **(D-F)** The expansion of CD8 and CD4 T-cell subsets on the bone marrow of mice treated using TMI/CSI2 vs. CSI2 alone. Representative flow cytometric analysis **(D)** and the percentages of specific T-cell subsets, such as naïve, central memory (CM) or effector memory/effector (EM/E) T-cells, together with the results of intracellular staining for IFNg **(E)**; means±SEM (n=3-6). **(F)** Reduced percentages of CMM cells (purple) and increased infiltration by CD8 T-cells (brown) in mice treated using using TMLI/CSI2 compared to CSI2 alone; means±SEM (n=3-6).

To assess the potential immunomodulatory effects of TMI/CSI2 treatment, we analyzed bone marrow-localized immune cell subsets within two weeks after TMI alone (4 Gy) and CSI2 alone (6 doses, 5mg/kg) or the combination. Flow cytometric analysis indicated activation of the total CD11b+ myeloid cells with elevated levels of MHC-II and CD86 costimulatory molecules (**Fig. 3B**), likely resulting from the antigen-presenting activity of macrophages (CD11b^+^F4/80^+^/MHC-II^HI^/CD86^+^) (**Fig. 3C**). In contrast, TMI/CSI2 treatment reduced the percentage of potentially tolerogenic, immature myeloid cells likely representing myeloid-derived suppressor cells (MDSC) (**Fig. 3C**). On the other hand, the combination treatment significantly increased both CD4+ and CD8^+^ T-cell populations in the BM compared to CSI2 alone (**Fig. 3D**) with specific expansion of central memory (CM) and effector memory/effector CD4 and CD8 T-cells compared to CSI2 alone (**Fig. 3E**). Intracellular cytokine staining indicated that TMI/CSI2 treatment dramatically induced the percentage of IFNγ-producing CD8 and CD4 T-cells in the BM (**Fig 3E**). These findings were further verified using immunohistochemical staining that showed increased CD8 T-cell infiltration and the reduced percentage of CMM leukemic cells in the BM of TMI/CSI2-treated mice compared to mice treated with CSI2 alone (**Fig 3F**).

Our previous studies on targeting AML using CpG-STAT3 inhibitors demonstrated that the immunotherapeutic effects of this strategy were not abrogated in TLR9KO mice. (38) (41) Thus, the effect of CSI seemed to primarily rely on TLR9-mediated immune activation of leukemic cells rather than the host immune cells. To determine whether TLR9 expression in the host is required for the efficacy of TMI/CSI2, we compared the effect of this strategy in TLR9 wild-type (WT) and knockout (KO) mice. As expected, TLR9-deletion in host immune cells did not impair the oligonucleotide uptake by CMM cells (**Supplementary Fig. S2)**.

In contrast to our previous studies in non-irradiated mice, TLR9KO mice showed reduced sensitivity to CSI2 combined with TMI (**Fig 4A-D**). Compared to WT mice, TLR9KO mice showed comparable reduction of CMM cells in the BM (**Fig 4E**) but the recruitment of CD8 T-cells (**Fig 4F**) and regulatory T-cells (**Fig 4G**) into spleen and BM in TLR9KO mice was significantly reduced. Thus, TLR9-mediated immune activation of host immune cells may at least partly contribute to the therapeutic effect of TMI/CSI2.

**Figure 4.**
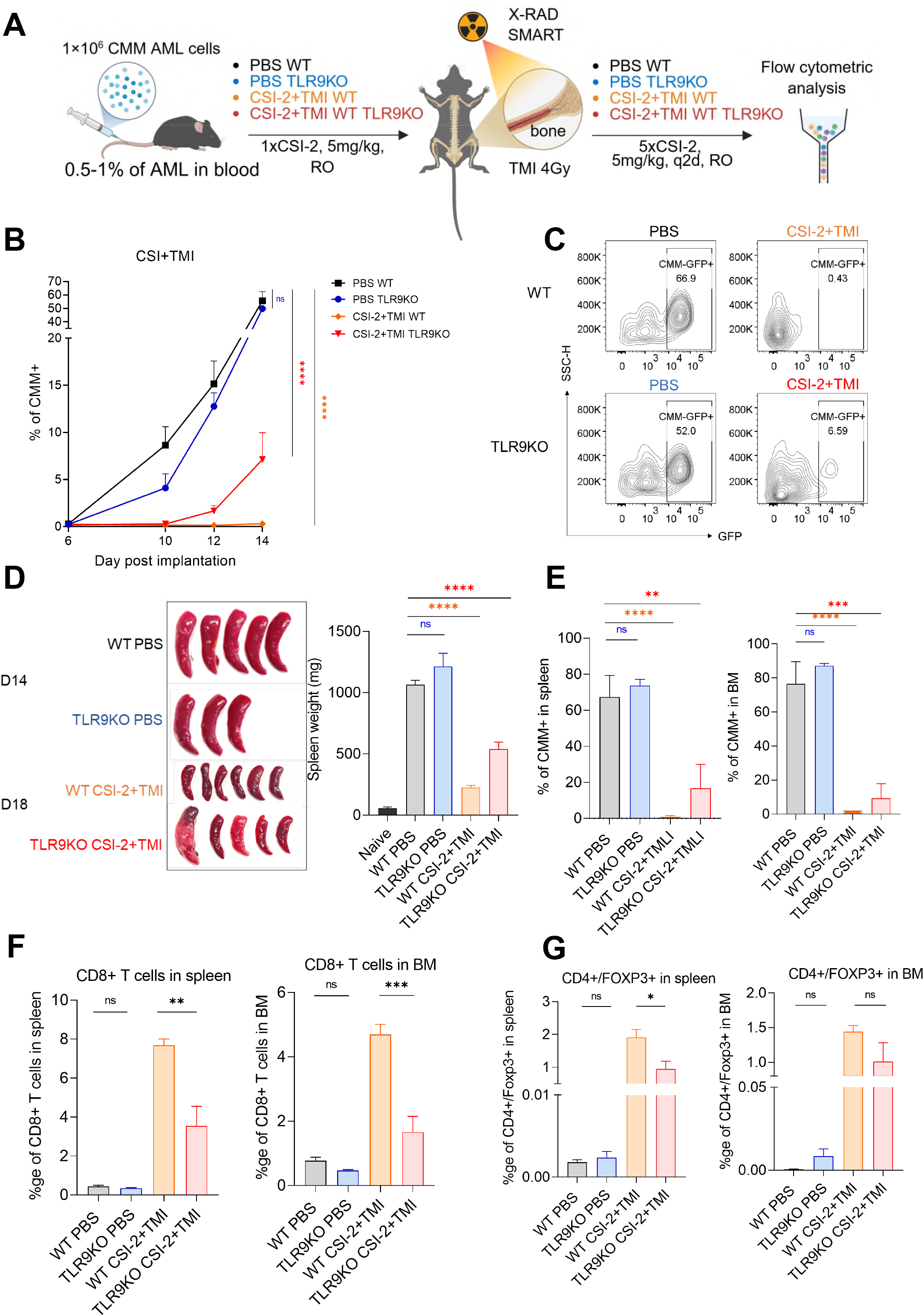
TLR9-deletion in host immune cells reduces but does not abrogate CD8 T-cell activation and anti-leukemic efficacy of TMI/CSI2 treatment. **(A)** C57BL/6 and TLR9KO/C57BL/6 mice were injected intravenously with 1×10^6^ CMM leukemia cells. Mice with established leukemia (0.5-1% of AML cells in blood) were treated 6 times daily with RO containing CSI2 (5mg/kg) or PBS. The first dose of treatment and/or without was combined with TMI irradiation at a dose of 4 (Gy) using Precision X-RAD SMART Plus/225cx (Precision X-Ray, CT, USA). **(B, C)** CMM leukemia progression monitored in blood using flow cytometric analysis. **(D)** Reduced spleen size (left) and weight (right) at the end of the study; **(E)** The percentages of CMM cells in spleen or BM as assessed using flow cytometry are not significantly different in TMI/CSI2-treated WT and TLR9KO mice. **(F)** The percentages of differentiating CD11b+ CMM leukemic cells in spleen or BM of TMI/CSI2-treated WT and TLR9KO mice. TLR9-deletion in the host reduces the recruitment of CD8+ T-cells **(G)** and Tregs (CD4+/Foxp3+) in spleen and bone marrow of TMI/CSI2-treated mice. Shown are means±SD, (n=3-6).

### TMI-CSI treated mice show sustained anti-tumor memory

As shown before, TMI-CSI-2 treatment improved survival in CMM-AML bearing mice compared to CSI-2 monotherapy. Notably, ∼20–30% of TMI-CSI-2–treated mice achieved complete disease-free survival, while the remaining 60–70% maintained only minimal disease (1–5% leukemic cells in peripheral blood; data not shown). Despite residual disease in some mice, all animals in the TMI-CSI-2 group survived for at least 12 additional months (a total of 15 months post-treatment), indicating the establishment of durable anti-tumor immune memory.

To directly test immune memory, we rechallenged TMI-CSI-2 and CSI-2 treated mice with CMM cells at 4–5-week post-initial therapy and monitored long-term survival. All TMI-CSI-2 treated mice remained disease-free beyond day 120 post-rechallenge, whereas only 20% of CSI-2 treated mice survived (**Fig 5**). These results demonstrate that TMI-CSI-2 therapy not only clears disease but also establishes robust, long-lasting immunological memory capable of rejecting secondary leukemic challenges.

**Figure 5.**
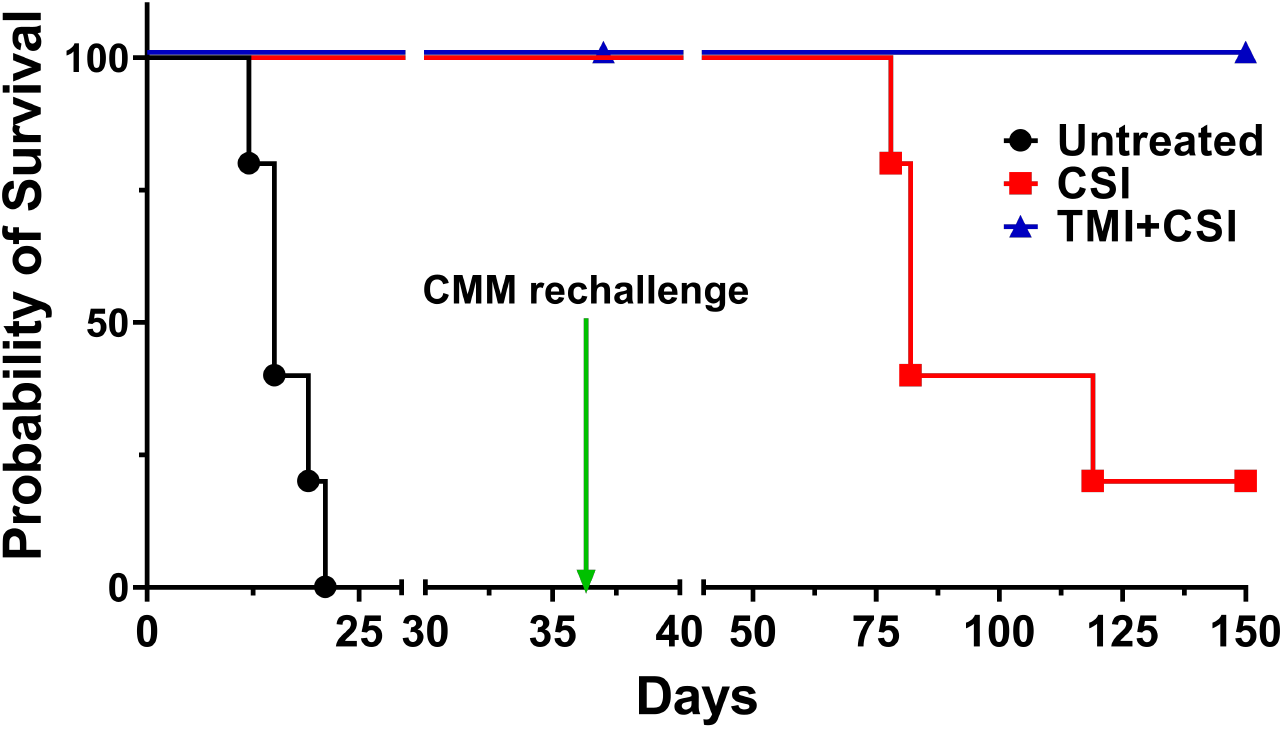
TMI/CSI2-treated mice generate immune resistance to CMM leukemia rechallenge. **(A)** CMM leukemia-bearing mice that survived after being treated using TMI/CSI2 or CSI2 alone were rechallenged using CMM cells 5–6-week later. All of the TMI/CSI2-treatd mice remained disease free for over 120 days. Survival was analyzed using Kaplan Meier Survival curve (Log Rank test; Mantel-cox).

## Discussion

Acute myeloid leukemia (AML) remains a clinically challenging disease, marked by poor long-term outcomes and high relapse rates, particularly in patients with advanced disease or limited eligibility for intensive therapies. While targeted agents and immunotherapies have begun to reshape the treatment landscape, there remains a critical need for combinatorial strategies that address both leukemic burden and immune escape. In this study, we present a novel therapeutic approach that combines targeted marrow irradiation (TMI) with STAT3 inhibition using the CpG-based decoy oligonucleotide CSI-2. This combination leverages the cytoreductive effects of organ-sparing radiation with the immunologic enhancement conferred by STAT3 blockade.

Radiotherapy is increasingly recognized for its immune-modulatory properties, including enhanced tumor immunogenicity, antigen presentation, and potential abscopal effects.[49–52] Further, we showed that this platform minimizes normal tissue toxicity compared to TBI-based approaches.[Ji Eun et al., 2022] Here, we demonstrate that TMI not only reduces leukemic burden but also promotes vascular remodeling in the bone marrow, improving blood flow and permeability. These changes enhance CSI-2 delivery and uptake in AML cells via TLR9 and scavenger receptor–mediated pathways.

Our prior work established that CSI-2 is an effective immunotherapeutic in AML models, particularly those involving the inv(16) subtype represented by CMM-AML.(41, 48) In moderate burden settings (∼1–5% CMM-GFP^+^ cells in peripheral blood and corresponding to 8-10% in BM), repeated CSI-2 treatment significantly reduced AML burden and eliminated leukemia-initiating cells, suggesting durable remission potential.(41, 49)] However, therapeutic effects of CSI-2 alone were mainly indirect and immune-mediated via CD8/CD4 T cells rather than directly cytotoxic. In immunodeficient NSG mice, lacking T, B and NK cell activity, the treatment had only limited effect on the same CMM leukemia.(41) Furthermore, in high disease burden models, the delayed kinetics of T cell priming (10–14 days) limits its efficacy, as mice often succumb to disease before immunity develops.

To overcome this limitation, we tested combination therapy with TMI and CSI-2. As expected, TMI alone (4 Gy) reduced leukemic burden at 48 hours post-treatment but conferred only a modest survival benefit (∼7–10 days over control). CSI-2 monotherapy improved median survival without significantly reducing disease burden by days 12–14. In contrast, the TMI-CSI-2 combination resulted in >80% survival at day 120, suggesting that neither cytoreduction nor immune activation alone is sufficient; rather, their synergy is required for sustained disease control in high-burden settings.

Mechanistically, we demonstrated that radiation enhances CSI-2 delivery to AML cells. TMI increased CSI-2 uptake in vivo, confirmed by both flow cytometry and quantitative multiphoton microscopy (QMPM).). In vitro, radiation upregulated TLR9 expression and CSI-2 uptake in CMM cells. This uptake was abrogated by co-treatment with CpG (TLR9 agonist) and dextran sulfate (scavenger receptor inhibitor), confirming a dual mechanism of CSI-2 internalization involving both receptors.

Previous work showed that CSI-2 retains efficacy in both WT and TLR9-deficient mice, indicating that TLR9 signaling in CMM cells—not in non-malignant immune cells—is the primary driver of antitumor immunity. Consistently, our TLR9-knockout mouse studies revealed that CSI-2 delivery and CD8^+^ T cell activation were only modestly reduced in the absence of host TLR9, supporting the notion that host TLR9 is dispensable for TMI-CSI-2–mediated immune responses.

TMI-CSI-2 treatment also enhanced T cell infiltration and activation within the BM. Compared to CSI-2 alone, the combination increased CD4^+^ and CD8^+^ T cells, particularly effector (CD44^+^CD62L^−^) and memory (CD44^+^CD62L^+^) subsets, and elevated IFNγ^+^ T cell frequencies.

Myeloid cells in the BM, including CD11b^+^F4/80^+^ macrophages and CD11b^+^Gr1^+^ neutrophils, upregulated MHC-II and CD86, indicating enhanced antigen-presenting function. Moreover, CMM cells themselves expressed co-stimulatory molecules (CD86/MHC-II) post-TMI-CSI-2 treatment, suggesting increased immunogenicity.

Notably, 20–30% of mice achieved disease-free survival, while the remaining 70–80% maintained minimal residual disease (1–5% CMM-GFP^+^ cells in peripheral blood) through days 90–120 (Data not shown) (n=5, survival study mice). Despite this, all mice survived >12 additional months, indicating long-term immune-mediated disease control. This was accompanied by increased effector memory T cells (CD44^+^CD62L^+^) in the BM. Rechallenge studies confirmed functional immune memory: all TMI-CSI-2–treated mice rejected AML upon re-injection, whereas most CSI-2 monotherapy mice failed to survive.

While TMI-CSI-2 showed robust efficacy in moderate-to-high disease burden (∼20–30%), its potential in even higher burden settings (>40%) warrants further evaluation. Importantly, TMI allows for selective dose escalation to skeletal and lymphoid tissues while sparing critical organs, prevent tissue damage in young and old mice.(13) In contrast, standard total body irradiation (TBI) at 8 Gy in combination with CSI-2 was highly toxic (median survival <30 days, with no disease at time of death), whereas TMI at 8 Gy with CSI-2 maintained >80% survival at 120 days (Data not shown). These results highlight the translational potential of TMI-CSI-2 as a safe and effective radio-immunotherapeutic strategy for AML, particularly in patients with high disease burden or relapse following standard therapies.

Our study demonstrates that combining targeted marrow irradiation (TMI) with STAT3 inhibition via CSI-2 produces a potent and durable anti-leukemic response in high-burden AML models. This approach leverages the direct cytoreductive effects of radiation and the immune-activating properties of STAT3 blockade to overcome disease persistence and immune tolerance. TMI enhances CSI-2 delivery, reprograms the bone marrow microenvironment, and supports robust T cell–mediated immunity and long-term immune memory. Its organ-sparing design and immunologic synergy make TMI-CSI-2 particularly promising for patients with relapsed or refractory AML—especially those with high STAT3 activity or who are ineligible for intensive therapies. These findings lay the foundation for future clinical translation of TMI-CSI-2 as a novel radio-immunotherapeutic platform.

## Authorship

Study design: S.S.M., S.H., M.K.; Data acquisition: Y-L.S, D.K., S.S.M., J-E.L., Data analysis and interpretation: Y-L.S., D.K., J-E.L.,S.S.M; Radiation treatment: D.Z, H.G, A.M.A, Intra vital microscopy imaging and image analysis: J.B, H.G. Administrative, technical, or material support: G.M., J.W., D.K., M.M., A.S., S.H., M.K.; Manuscript writing: S.S.M., S.H., M.K.

## Acknowledgements

We acknowledge dedication of staff members of the Analytical Cytometry, Analytical Pharmacology, DNA/RNA Synthesis, Integrative Genomics/Bioinformatics, Light Microscopy/Digital Imaging, Research Pathology, Small Animal Imaging and Animal Resource Core Laboratories supported by the NCI/NIH grant number P30CA033572. The content is solely the responsibility of the authors and does not necessarily represent the official views of the NIH.

