## Supplementary Figures for "Targeted Irradiation and STAT3 Inhibition Reprogram the AML Microenvironment and Extend Survival: Toward Translational Immunoradiotherapy"

### Supplemental figures

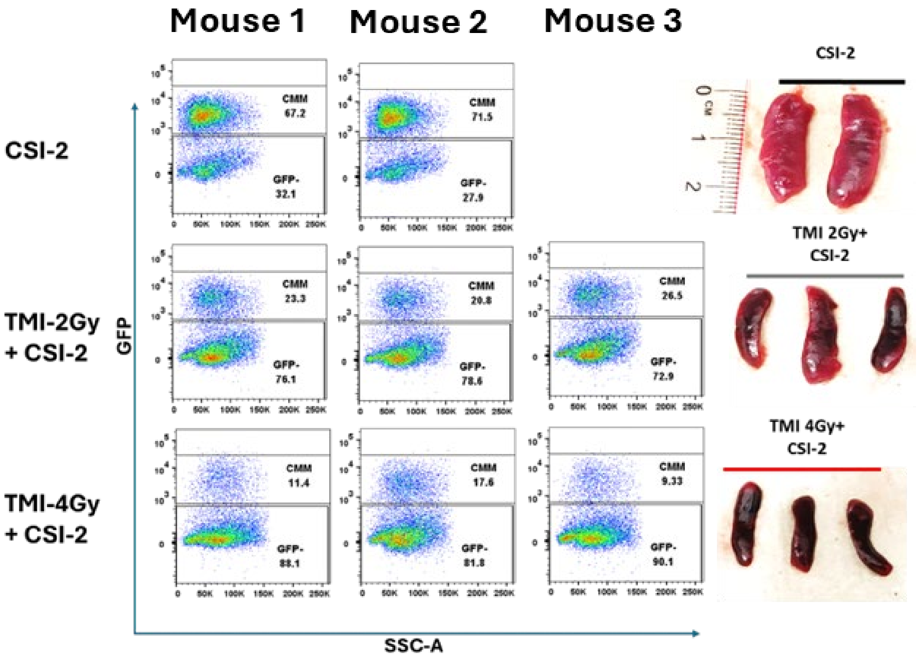


**Supplemental Fig S1: Dose-dependent effect of TMI on the AML burden in mice.** Leukemia cell killing was higher in 4 Gy than 2 Gy as shown by reduced CMM-GFP+ cells in BM by flow cytometry and reduced spleen size.

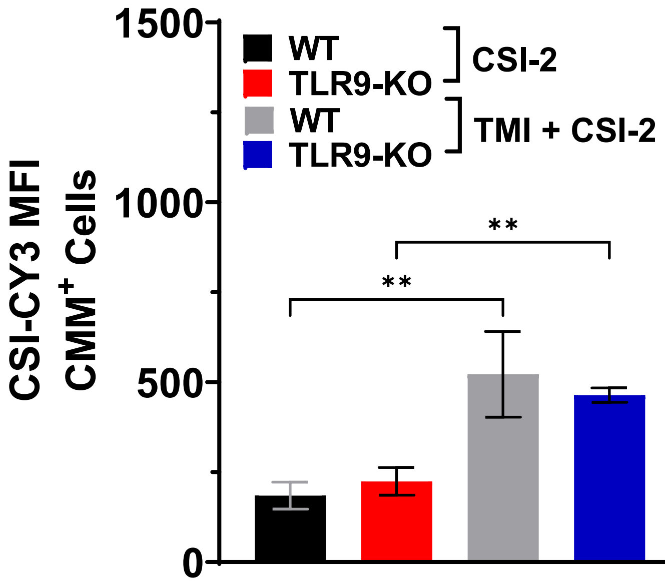


**Supplemental Fig S2: Host TLR9 cells was indispensable for CSI-2 uptake in CMM cells.** CSI-$2^{Cy3}$ uptake in WT and TLR9-KO mice was not affected after TMI.
